# Loss of *Nupr1* promotes engraftment by tuning the quiescence threshold of hematopoietic stem cell repository via regulating p53-checkpoint pathway

**DOI:** 10.1101/2020.07.16.205898

**Authors:** Tongjie Wang, Chengxiang Xia, Qitong Weng, Kaitao Wang, Yong Dong, Sha Hao, Fang Dong, Xiaofei Liu, Lijuan Liu, Yang Geng, Yuxian Guan, Juan Du, Tao Cheng, Hui Cheng, Jinyong Wang

## Abstract

Hematopoietic stem cells (HSCs) are dominantly quiescent under homeostasis, which is a key mechanism of maintaining the HSC pool for life-long hematopoiesis. Dormant HSCs poise to be immediately activated on urgent conditions and can return to quiescence after regaining homeostasis. To date, the molecular networks of regulating the threshold of HSC dormancy, if exist, remain largely unknown. Here, we unveiled that deletion of *Nupr1*, a gene preferentially expressed in HSCs, activated the quiescence HSCs under homeostatic status, which conferred engraftment competitive advantage on HSCs without compromising their stemness and multi-lineage differentiation abilities in serial transplantation settings. Following an expansion protocol, the *Nupr1*^*-/-*^ HSCs proliferate more robustly than their wild type counterparts *in vitro. Nupr1* inhibits the expression of p53 and the rescue of which offsets the engraftment advantage. Our data unveil the *de novo* role of *Nupr1* as an HSC quiescence-regulator, which provides insights into accelerating the engraftment efficacy of HSC transplantation by targeting the HSC quiescence-controlling network.

## Introduction

Hematopoietic stem cells (HSCs), the seeds of adult blood system, generate all the blood lineages via hierarchical hematopoiesis. Under steady-state, the majority of HSCs are maintained in quiescence to reserve the HSC pool for life-long hematopoiesis^1^. However, the dormant HSCs can be rapidly activated for stress hematopoiesis on emergency conditions, such as excessive blood loss, radiation injury, and chemotherapy damage^2^. Mounting evidence point to the existence of intrinsic molecular machinery of regulating HSC dormancy. In haploinsufficient *Gata2*^+/-^ mice, HSCs show mildly increase of quiescent cells on homeostasis condition^3^. Dnmt3a-knockout (KO) hematopoietic stem cells (HSCs) had increased self-renewal ability and expanding HSC numbers in the bone marrow^4, 5^. JunB inactivation deregulates the cell-cycle machinery and reduces quiescent HSCs^6^. *Hif-1α*-deficient HSCs also show decreased dormant HSCs^7^. Conditional knockout of cylindromatosis (CYLD) induced dormant HSCs exit quiescence and abrogated their repopulation and self-renewal potential^8^. CDK6, a protein not expressed in long-term HSCs but short-term HSCs, regulates the quiescence exit in human hematopoietic stem cells, and overexpression of which promotes engraftment^9^. To date, the underlying signaling regulatory network of HSC quiescence remains largely unknown.

NUPR1 (Nuclear protein transcription regulator 1) is a member of the high-mobility group of proteins, which was first discovered in the rat pancreas during the acute phase of pancreatitis and was initially called p8^10^. The same gene was discovered in breast cancer and was named as Com1^11^. NUPR1 demonstrates various roles involving apoptosis, stress response, and cancer progression, which depends on distinct cellular context. In certain cancers, such as breast cancer, NUPR1 inhibits tumor cell apoptosis, induces tumor establishment and progression^12-15^. On the contrary, in prostate cancer and pancreatic cancer, NUPR1 shows tumor-growth inhibitory effect^16, 17^. Accumulated studies reveal that NUPR1 is a stress-induced protein: interference of NUPR1 can upregulate the sensitivity astrocytes to oxidative stress^18^; loss of it can promote resistance of fibroblasts to adriamycin-induced apoptosis^19^; NUPR1 mediates cannabinoid-induced apoptosis of tumor cells^20^; overexpression of *NUPR1* can negatively regulate MSL1-dependent HAT activity in Hela cells, which induces chromatin remodeling and relaxation allowing access to DNA of the repair machinery^21^. Nonetheless, the potential roles of *Nupr1*, which is preferentially expressed in HSCs among the HSPC, in hematopoiesis remain elusive. NUPR1 interacts with p53 to regulate cell cycle and apoptosis responding to stress in breast epithelial cells^19, 22^. p53 plays several roles in homeostasis, proliferation, stress, apoptosis, and aging of hematopoietic cells^23-27^. Deletion of p53 upregulates HSC self-renewal but impairs their repopulating ability and leads to tumors^28^. Hyperactive expression of p53 in HSCs decreased the HSC pool size, reduced engraftment and deep quiescence^29-31^. These reports support the essential check-point role of p53 in regulating HSC fate. Nonetheless, it is unknown whether NUPR1 and p53 coordinately regulate the quiescence of HSCs.

Here, we used a *Nupr1* conditional knockout model to investigate the consequences of loss of function of *Nupr1* in HSC context. *Nupr1*-deletion in HSCs led to their quiescence withdrawal under homeostasis. In a competitive repopulation setting, *Nupr1*-deleted HSCs robustly proliferated and showed dominant engraftment over wild type counterparts. Besides, *Nupr1*-deleted HSCs expanded abundantly and preserved their stemness in vitro in comparison with wild type HSCs. The rescued expression of p53 by *Mdm2*^+/-^ offset the effects introduced by loss of *Nupr1* in HSCs. Our studies reveal the *de novo* role and signaling mechanism of *Nupr1* in regulating the quiescence of HSCs.

## Methods

### Mice

Animals were housed in the animal facility of the Guangzhou Institutes of Biomedicine and Health (GIBH). *Nupr1*^fl/fl^ mice were constructed by Beijing Biocytogen Co., Ltd. CD45.1, Vav-cre, Mx1-cre, and *Mdm2*^+/-^ mice were purchased from the Jackson laboratory. All the mouse lines were maintained on a pure C57BL/6 genetic background. All experiments were conducted in accordance with experimental protocols approved by the Animal Ethics Committee of GIBH.

### HSC cell cycle analysis

We first labeled the HSCs with (CD2, CD3, CD4, CD8, Ter119, B220, Gr1, CD48)-Alexa Flour700, Sca1-Percp-cy5.5, c-kit-APC-cy7, CD150-PE-cy7, CD34-FITC and CD135-PE. Then the cells were fixed using 4% PFA. After washing, the fixed cells were permeabilized with 0.1% saponin in PBS together with the Ki-67-APC staining for 45 minutes. Finally, the cells were resuspended in DAPI solution for staining 1 hour. The data were analyzed using Flowjo software (FlowJo).

### BrdU incorporation assay

*Nupr1*^-/-^ mice and WT littermate mice were injected with 1 mg BrdU on Day 0. Then they were fed with water containing BrdU (0.8 mg/mL). On Day 3, 4, 5 after the injection of BrdU, four mice of each group were sacrificed. The incorporation rates of BrdU were analyzed by flow cytometry according to the BD Pharmingen (tm) APC BrdU Flow Kit instructions.

### HSC culture

The HSC culture protocol is as described ^32^. Briefly, fifty HSCs were sorted into fibronectin (Sigma)-coated 96-well U-bottom plate directly and were cultured in medium composed of F12 medium (Life Technologies), 1% insulin–transferrin–selenium–ethanolamine (ITSX; Life Technologies), 10 mM HEPES (Life Technologies), 1% penicillin/streptomycin/glutamine (P/S/G; Life Technologies), 100 ng/ml mouse TPO, 10 ng/ml mouse SCF and 0.1% PVA (P8136). Complete medium changes were made every 2–3 days, by manually removing medium by pipetting and replacing fresh medium as indicated.

### Limiting dilution assay

For limiting dilution assays^33^, the 10-day cultured cells were transplanted into lethally irradiated C57BL/6-CD45.1 recipient mice, together with 2×10^5^ CD45.1 bone-marrow competitor cells. Donor chimeras was analyzed as above. Limiting dilution analysis was performed using ELDA software^34^, based on a 1% peripheral-blood multilineage chimerism as the threshold for positive engraftment.

### Bone marrow competitive repopulation assay

One day before bone marrow transplantation, adult C57BL/6 (CD45.1, 8-10 weeks old) recipient mice were irradiated with 2 doses of 4.5Gy (RS 2000, Rad Source) for a 4-hour interval. Two hundred and fifty thousand BMNCs from *Nupr1*^-/-^ mice (CD45.2) and equivalent WT (CD45.1) counterparts were mixed and injected into irradiated CD45.1 recipients by the retro-orbital injection. Control BMNCs (CD45.2), *Mdm2*^+/-^*Nupr1*^-/-^ BMNCs (CD45.2) or *Mdm2*^*+/-*^ BMNCs (CD45.2) were also mixed with equivalent competitors (CD45.1) and transplanted into recipients. The transplanted mice were maintained on trimethoprim-sulfamethoxazole-treated water for 2 weeks. For secondary transplantation, BMNCs of primary competitive transplanted recipients were obtained. One million of total BMNCs were injected into irradiated CD45.1 recipients (2 doses of 4.5Gy, one day before transplantation). Donor-derived cells and hematopoietic lineages in PB were assessed monthly by flow cytometry.

## Results

### Loss of *Nupr1* accelerates the turn-over rates of HSCs under homeostasis

A majority of long-term HSCs are quiescent under homeostasis, which is a key mechanism for maintaining the HSC pool for life-long steady hematopoiesis. We hypothesize that among those genes, preferentially expressed in HSCs but immediately down regulated in MPPs, might form an intrinsic regulatory network for maintaining the HSC quiescence. To test our hypothesis, we explored such factor candidates by RNA-Seq analysis of the sorted HSCs (Hematopoietic stem cells, Lin^-^ CD48^-^ Sca1^+^ c-kit^+^ CD150^+^) and MPPs (Multipotent stem cells, Lin^-^ Sca1^+^ c-kit^+^ CD150^-^). Differential expression gene analysis showed a pattern of HSC-preferential transcription factors, including *Rorc, Hoxb5, Rarb, Gfi1b, Mllt3*, and *Nupr1*. By literature search, we found that most of the candidate genes were reportedly not involved in regulating HSC homeostasis. Thus, we focus on the *Nupr1* gene, the role of which in hematopoiesis has not been reported. The expression of *Nupr1* in HSCs is significantly higher (> 25-fold, p = 0.002) than MPPs (Figure 1A, left). Real-Time PCR further confirmed the same expression pattern (p <0.001), implicating an unknown role of *Nupr1* in HSCs (Figure 1A, right).

**Fig 1.**
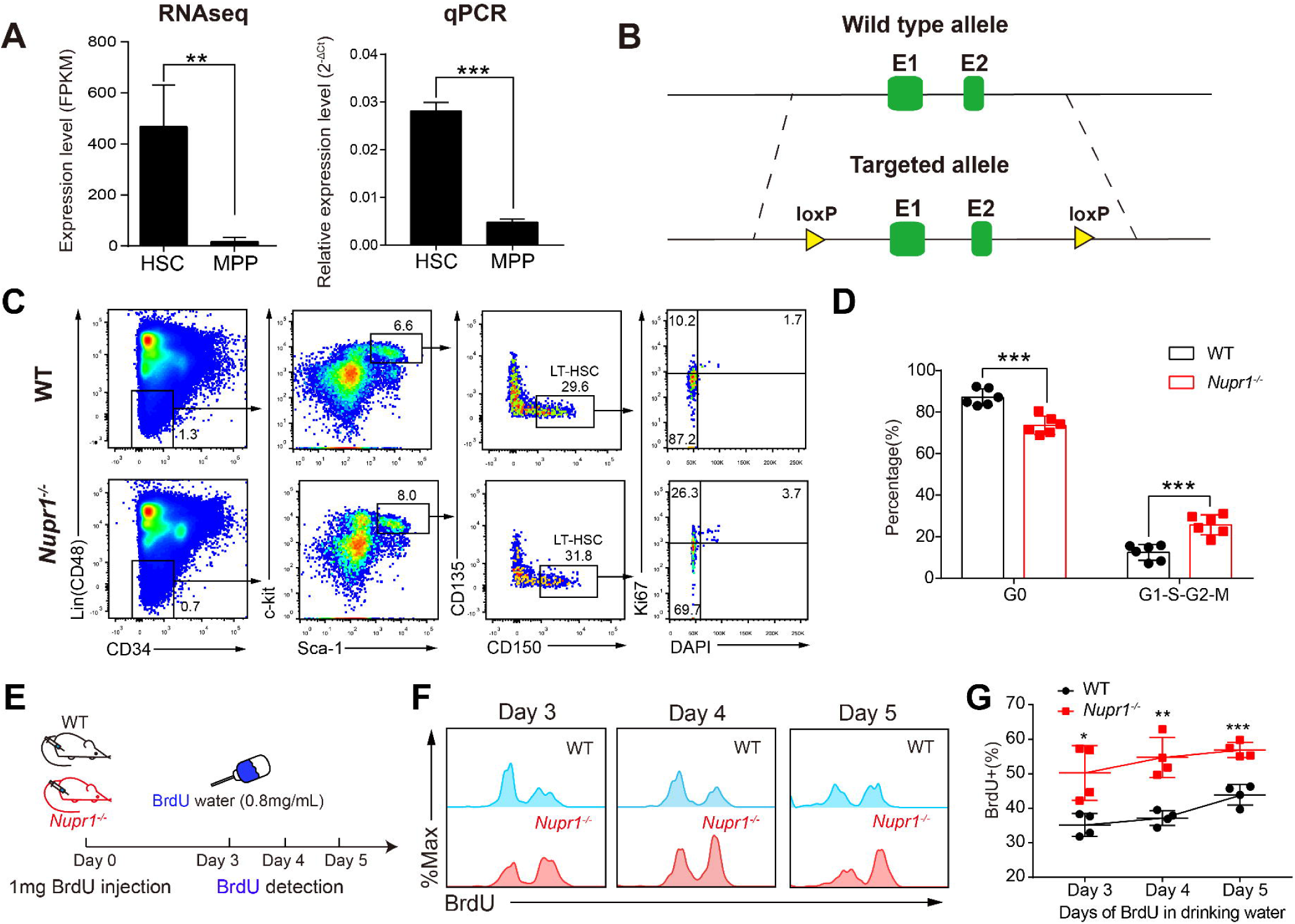
Loss of *Nupr1* activates dormant HSCs under homeostasis. (A) Expression pattern of *Nupr1* in hematopoietic stem cells (HSCs) and multipotent progenitors (MPPs) examined by RNA-sequencing and Real-Time PCR. One thousand HSC or MPP cells from bone marrow of wild type mice were sorted as individual samples for RNA-sequencing (n=4). HSCs are defined as Lin (CD2, CD3, CD4, CD8, Mac1, Gr1, Ter119, B220)^-^, CD48^-^, Sca1^+^, c-kit^+^, and CD150^+^. MPPs are defined as Lin (CD2, CD3, CD4, CD8, Mac1, Gr1, Ter119, B220)^-^, Sca1^+^, c-kit^+^, and CD150^-^. Data are analyzed by unpaired Student’s t-test (two-tailed). **p < 0.01, ***p<0.001. Data are represented as mean ± SD (qPCR, n = 3 mice for each group). (B) Targeting strategy of knockout of *Nupr1* gene in mouse. Wild type *Nupr1* exons 1, and 2 are shown as green boxes. Two loxp elements flanking exon 1 and exon 2 were inserted. (C) Cell cycle analysis of *Nupr1*^-/-^ HSCs under homeostasis. Representative plots of cell cycle from representative WT and *Nupr1*^-/-^ mice (8-week-old). WT littermates (8-week-old) were used as control. HSCs (Lin^-^ (CD2^-^ CD3^-^ CD4^-^ CD8^-^ B220^-^ Gr1^-^ CD11b^-^ Ter119^-^) CD48^-^ Sca1^+^ c-kit^+^ CD150^+^ CD34^-^ CD135^-^) were analyzed by DNA content (DAPI) versus Ki-67. G0 (Ki-67^low^DAPI^2N^), G1 (Ki-67^high^DAPI^2N^), G2-S-M (Ki-67^high^DAPI^>2N-4N^). (D) Statistical analysis of HSC cell cycle. The percentages (%) of HSCs in G0, G1-G2-S-M stages were analyzed. Data are analyzed by unpaired Student’s t-test (two-tailed). **p < 0.01. Data are analyzed by Student’s t-test. Data are represented as mean ± SD (n = 6 mice for each group). (E) The strategy of BrdU incorporation assay. The 8-week-old *Nupr1*^-/-^ mice and littermates were injected intraperitoneally with 1mg BrdU on day 0. Then the mice were continuously fed with BrdU (0.8mg/ml) water until analyzed on day 3, 4, and 5. (F) Dynamic tendency analysis of BrdU^+^ HSCs after BrdU administration by flow cytometry on day 3, 4, and 5. (G) Ratio kinetics of BrdU^+^ HSCs. Data are analyzed by unpaired Student’s t-test (two-tailed). *p < 0.05, **p < 0.01, ***p < 0.001. Data are analyzed by Student’s t-test. Data are represented as mean ± SD (n = 4 mice for each group).

To study whether *Nupr1* has any potential impact on the hematopoiesis of HSCs, we constructed the *Nupr1* conditional knockout mice by introducing two loxp elements flanking the exon 1 and 2 of *Nupr1* locus using a C57BL/6 background mESC line (Figure 1B). The generated *Nupr1*^fl/fl^ mice were further crossed to Vav-Cre mice to generate *Nupr1*^fl/fl^; Vav-Cre compound mice (*Nupr1*^-/-^ mice). The deletion of *Nupr1* was confirmed by PCR validation of genotype and qPCR of *Nupr1* expression level (Supplementary Figure 1). Adult *Nupr1*^-/-^ mice (8-10-week-old) had a normal percentage of blood lineage cells in peripheral blood, including CD11b^+^ myeloid, CD19^+^ B, and CD3^+^ T lineage cells (Supplementary Figure 2). We further investigated the potential alterations of HSC hemostasis in the absence of *Nupr1*. Flow cytometry analysis demonstrated that *Nupr1*^-/-^ HSC pool was comparable to wild type counterparts in terms of ratios and absolute numbers (Supplementary Figure 3). Subsequently, we examined the cell cycle status of *Nupr1*^-/-^ HSCs using the proliferation marker Ki-67 and DAPI staining and found that the ratio of *Nupr1*^-/-^ HSCs in G0-status was reduced significantly (p<0.001). Compared with those of WT HSCs (median value: *Nupr1*^-/-^ HSCs =73.67%, WT HSCs = 87.15%), more *Nupr1*^-/-^ HSCs entered G1-S-S2 and M phase (Figure 1C, D). To further confirm this novel phenotype, we performed BrdU incorporation assay, which is conventionally used for assessing the turn-over rates of blood cells *in vivo*^35^. The 8-week-old *Nupr1*^-/-^ mice and littermates were injected intraperitoneally with 1mg BrdU on day 0, followed by administration of BrdU via water feeding (0.8 mg/ml) for up to 5 days (Figure 1E). After three days of BrdU labeling, ∼50% of *Nupr1*^-/-^ HSCs became BrdU^+^ compared with ∼35% of WT HSCs. Kinetic analysis with BrdU incorporation from day 3 to day 5 revealed that *Nupr1*^-/-^ HSCs contained a 1.5-fold higher BrdU^+^ population over WT HSCs (Figure 1F, G). Collectively, these data indicate that the *Nupr1*-deletion drives HSCs entering cell cycle and accelerates their turn-over rates on homeostasis.

### *Nupr1*^-/-^ HSCs show repopulating advantage without compromising multi-lineage differentiation capacity

To confirm whether *Nupr1*^-/-^ HSCs have repopulating advantage or disadvantage in *vivo*, we performed typical HSC-competitive repopulation assay. Two hundred and fifty thousand whole bone marrow nucleated cells (BMNCs) from *Nupr1*^-/-^ mice (CD45.2) were transplanted into lethally irradiated recipients (CD45.1) along with equivalent WT (CD45.1) competitors. BM cells from the littermate mice (Vav-Cre^+^, CD45.2^+^) were mixed with WT (CD45.1) competitors and transplanted into the recipients as the experiment control. Sixteen weeks later, one million BMNCs of the primary recipients were transplanted into lethally irradiated recipients for assessing long-term engraftment (Figure 2A). We observed that donor *Nupr1*^-/-^ cells took about ∼70% in the primary recipients, while the control cells accounted for 50%-60% in the recipients. *Nupr1*^-/-^ cells gradually dominated in peripheral blood of recipients over time after transplantation (Figure 2B). In the chimeras, ∼70% of myeloid cells and B lymphocytes were *Nupr1*^-/-^ donor-derived cells, while ∼60% of T lymphocytes were CD45.1 competitive cells (Figure 2C). To further explore whether *Nupr1*^-/-^ HSCs dominate in chimer, we sacrificed the chimeras and analyzed the HSCs 16 weeks after transplantation. Compared with the control group, the proportion and absolute number of *Nupr1*^-/-^ HSCs were significantly more (∼1.5 folds) than the control HSC in primary recipients (Figure 2D, E). Previous research reported that HSCs proliferated rapidly at the expense of their long-term repopulating ability^36-40^. Interestingly, consistent with the dominating trend in primary transplantation, *Nupr1*^-/-^ cells continuously dominated in secondary recipients (Figure 3A). *Nupr1*^-/-^ HSCs occupied up to 90% of the total HSCs in the bone marrow (BM) of secondary recipients. However, the control HSCs were less than 10% in the secondary recipients (Figure 3B, C). In aggregate, these results indicate that the deletion of *Nupr1* promotes the repopulating ability of HSCs without impairing their long-term engraftment ability.

**Fig 2.**
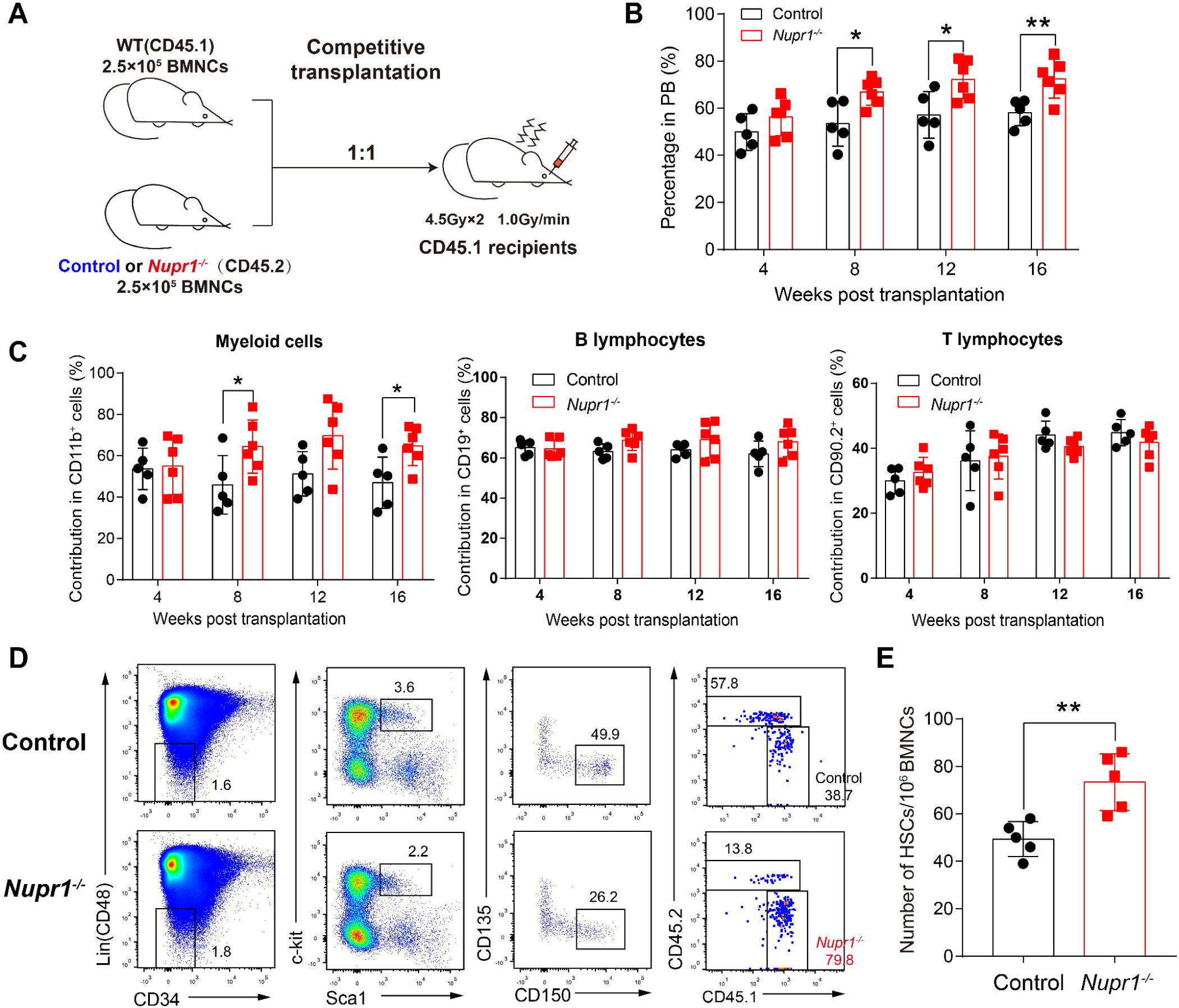
*Nupr1*^-/-^ HSCs show repopulating advantage in competitive transplantation. (A) Schematic diagram of competitive transplantation assay. 2.5 × 10^5^ *Nupr1*^-/-^ BMNCs (CD45.2) or littermate control BMNCs (Vav-cre^+^, CD45.2) were mixed with equivalent WT (CD45.1) counterparts and injected into individual lethally irradiated recipients (CD45.1). Four months later, the recipients were sacrificed. One million BMNCs from primary transplanted recipients were transplanted to lethally irradiated secondary recipients. (B) Kinetic analysis of donor chimeras (CD45.2^+^) in peripheral blood. Data are analyzed by paired Student’s t-test (two-tailed). *p<0.05, **p < 0.01. Data are analyzed by Student’s t-test. Data are represented as mean ± SD (Control group: n = 5 mice, *Nupr1*^*-/-*^ group: n =6 mice). (C) Kinetic analysis of donor-derived lineage chimeras in peripheral blood, including myeloid cells (CD11b^+^) (left), B lymphocytes (CD19^+^) (middle), and T lymphocytes (CD90.2^+^) (right) in peripheral blood. Data are analyzed by paired Student’s t-test (two-tailed). *p < 0.05. Data are analyzed by Student’s t-test. Data are represented as mean ± SD (Control group: n = 5 mice, *Nupr1*^*-/-*^ group: n = 6 mice). (D) Flow cytometry analysis of HSC compartment in primary recipients four months after transplantation. Representative plots from one recipient mouse in each group are shown. (E) Cell number of donor-derived HSCs in primary recipients four months after competitive transplantation. Data are analyzed by Student’s t-test. **p < 0.01. Data are represented as mean ± SD (n = 5).

**Fig 3.**
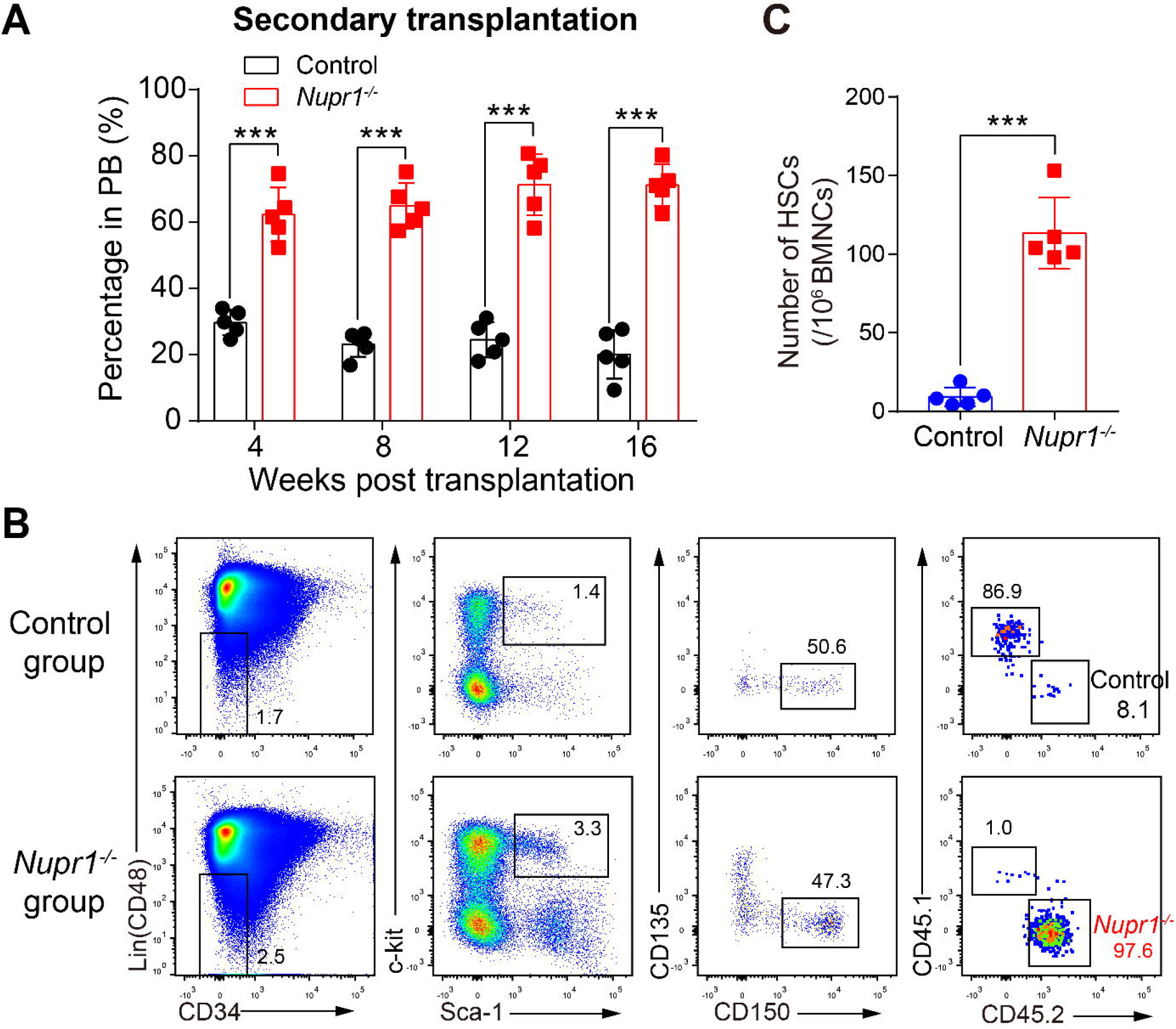
*Nupr1*^-/-^ HSCs continuously show competitive advantage without losing their long-term self-renew ability in secondary transplantation. (A) Kinetic analysis of donor chimeras (CD45.2^+^) in peripheral blood of secondary transplanted recipients. Data are analyzed by paired Student’s t-test (two-tailed). ***p < 0.001. Data are represented as mean ± SD (n = 5 mice). (B) Flow cytometry analysis of donor *Nupr1*^-/-^ HSCs in secondary recipients 16 weeks after transplantation. Representative plots from each group mice were shown. (C) Cell number of donor-derived HSCs in secondary recipients four months after competitive transplantation. Data are analyzed by unpaired Student’s t-test (two-tailed). ***p < 0.001. Data are represented as mean ± SD (n= 5 mice).

### *Nupr1*^-/-^ hematopoietic stem cells are highly sensitive to irradiation-stress but re-cover fast

HSCs under cell cycle were reported to be more sensitive to irradiation-damage ^41^. To explore whether *Nupr1*^*-/-*^ HSCs with faster turn-over rate are more sensitive to irradiation, WT mice and *Nupr1*^*-/-*^ mice were exposed to a single dose of total body irradiation (4Gy dose, 1Gy/min). Then we analyzed the apoptosis and cell cycle status 6 hours (early stage) and 24 hours (late stage) later. As expected, *Nupr1*^*-/-*^ HSCs showed a significantly enhanced sensitivity to irradiation: only ∼40% of *Nupr1*^*-/-*^ HSCs lived 6 hours after irradiation, whereas ∼70% of WT HSCs were still alive (Supplementary Figure 4A). The proportion of radiation-induced apoptosis (Annexin V^+^) of *Nupr1*^*-/-*^ HSCs was significantly (p < 0.05) higher (∼ 2-fold) than that of WT HSCs (Supplementary Figure 4A). Furthermore, ∼60% of the residual *Nupr1*^*-/-*^ HSCs were in G1-S-G2-M proliferative phases compared with ∼50% of the residual WT HSCs (p < 0.05, Supplementary Figure 4B), indicating an accelerated replenish rate in response to irradiation damage. At a later stage (24 hours) after irradiation, we observed more lived *Nupr1*^*-/-*^ HSCs (WT vs. *Nupr1*^*-/-*^: 74% vs. 86%) and less irradiation-induced apoptotic *Nupr1*^*-/-*^ HSCs compared with WT HSCs (Supplementary Figure 4C). The proportion of cycling cells (G1-S-G2-M proliferative phases) in the residual *Nupr1*^*-/-*^ HSCs was still significantly (p < 0.001) higher than the WT HSCs 24 hours after irradiation (WT vs. *Nupr1*^*-/-*^: 29% vs. 48%) (Supplementary Figure 4D). Taken together, *Nupr1*^*-/-*^ HSCs are susceptible to the irradiation-induced damage, but the survived HSCs recovered more accelerating turn-over rates.

### *Nupr1*-deleted HSCs expand robustly *in vitro*

We next examined whether the deletion of *Nupr1* could enhance HSC expansion *in vitro*. Fifty HSCs sorted from WT and *Nupr1*^-/-^ mice were cultured *in vitro* for 10 days as previously described^32^ (Figure 4A). After 10-day-culture, the wild type input cells achieved a yield of more than 2 × 10^4^ cells, while *Nupr1*^-/-^ HSCs produced approximately 5×10^4^ total cells (p <0.001, Figure 4B). The colonies derived from *Nupr1*^-/-^ HSCs were much larger than WT HSCs (Figure 4C). Furthermore, we analyzed the phenotypic HSC populations in the expanded cells and found that the absolute number of phenotypic HSC in individual *Nupr1*^-/-^ colonies were 3 times more than WT HSCs (p=0.005, Figure 4D, E). To determine whether the quantitative expansion of phenotypic HSCs contains net proliferation of functional HSCs, we performed competitive repopulating unit (CRU) assays^33^, using the serial doses of limiting dilutions of the in vitro expanded cells. The WT HSC frequency in the 10-day expanded cells is 1 in 371 cells, which is equivalent to 61 functional HSCs. While the *Nupr1*^-/-^ HSC frequency in the 10-day expanded cells is 1 in 190 cells (Figure 4F)^34^, which is equivalent to 263 functional HSCs (p=0.045). Therefore, the deletion of *Nupr1* induced around four-fold expansion in functional HSC number over the WT HSCs. Deletion of *Nupr1* enhances the expansion ability of HSCs *in vitro*.

**Fig 4.**
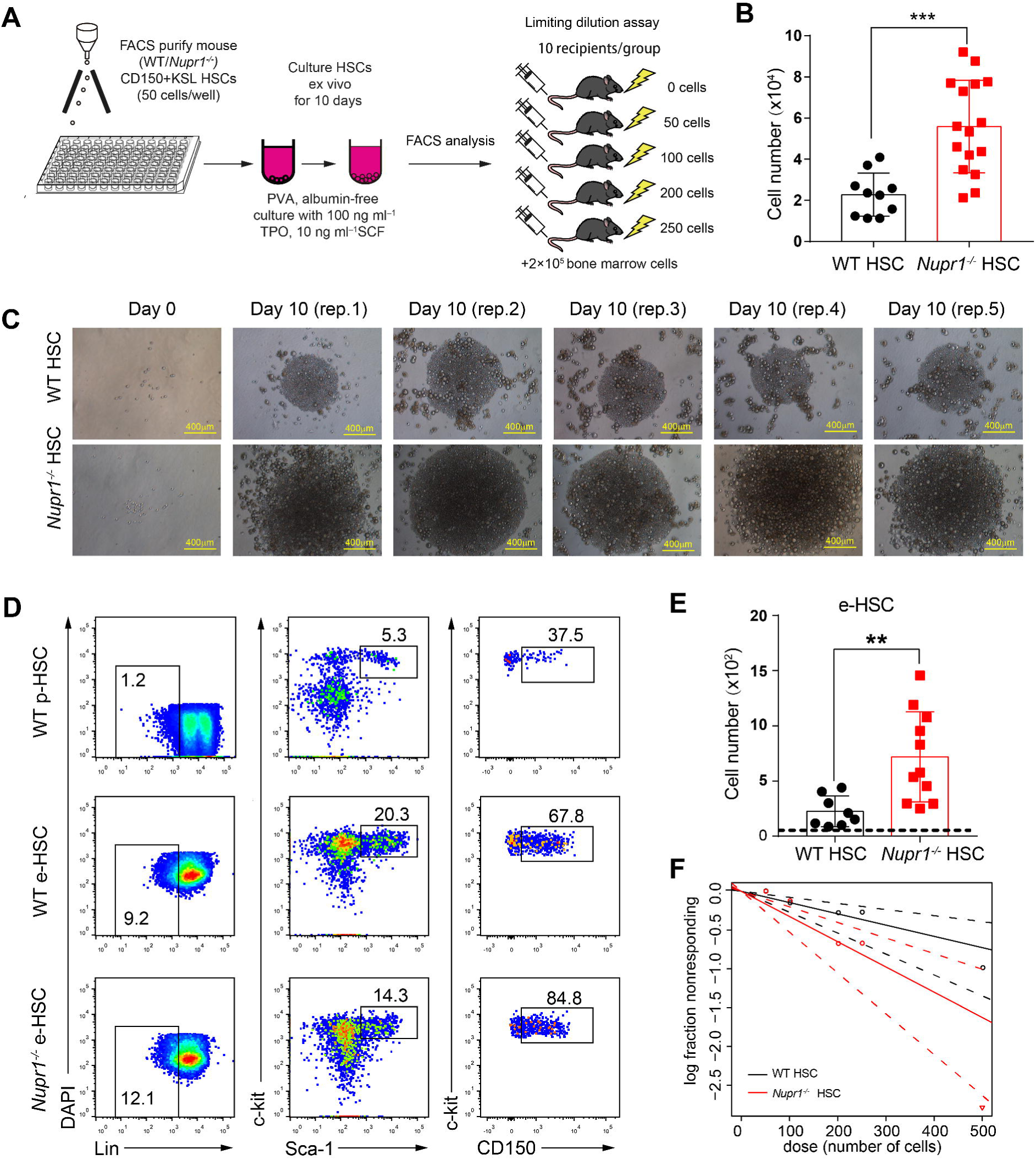
Deletion of *Nupr1* promotes HSC expansion in vitro. (A) Schematic diagram of the HSC expansion in vitro. 50 CD150^+^KSL HSCs (from WT and *Nupr1*^*-/-*^ mice) were sorted into fibronectin-coated plate wells, containing albumin-free F12 medium supplemented with 1 mg/ml PVA, 100 ng/ml TPO and 10 ng/ml SCF. HSCs were cultured for 10 days and then analyzed by flow cytometry. For limiting dilution assay, serial doses were transplanted into lethally irradiated recipients, together with 2×10^5^ bone-marrow competitor cells. (B) Cell number derived from 50 HSCs after a 10-day-long culture in *vitro*. Data are analyzed by unpaired Student’s t-test (two-tailed). ***p < 0.001. Data are represented as mean ± SD (WT, n = 10; *Nupr1*^*-/-*^, n=16) (C) Representative images of WT and *Nupr1*^*-/-*^ HSCs from freshly isolated HSCs (Day 0) and 10-day-long cultures (Day 10). Images of five representative colonies (biological replicates) are shown. (D) Representative plots of HSC analysis by flow cytometry from cultured WT and *Nupr1*^*-/-*^ HSCs at day 10. p-HSC indicates primary HSCs from BM. e-HSC indicates expanded HSCs after 10-day culture *ex vivo*. (E) Cell counts of phenotypic CD150^+^KSL HSCs at day 10 after culture. The dashed indicates the primary input cell amount. Data are analyzed by unpaired Student’s t-test (two-tailed). **p < 0.01. Data are represented as mean ± SD (WT, n = 8; *Nupr1*^*-/-*^, n=11). (F) Poisson statistical analysis after limiting-dilution analysis; plots were obtained to allow estimation of CRU content within each condition (n = 10 mice transplanted at each dose per condition, *p<0.05). The plot shows the percentage of recipient mice containing less than 1% CD45.2^+^ cells in the peripheral blood at 16 weeks after transplantation versus the number of cells injected per mouse. *p<0.05.

### Reversion of p53 expression offsets the competitiveness of *Nupr1*^-/-^ HSCs

To further investigate the underlying molecular mechanisms of *Nupr1* in regulating HSCs, we performed RNA-Seq analysis of *Nupr1*^-/-^ HSCs from 8-week-old *Nupr1*^-/-^ mice. Differential gene-expression (DEGs) analysis indicated that there were 319 differential genes exists between the WT and *Nupr1*^*-/-*^ HSCs (a difference in expression of over 2-fold; adjusted P value < 0.05 (DESeq2 R package)). Gene-ontology analysis of these differentially expressed genes indicated enrichment for genes involved in regulation of mitotic cell cycle and negative regulation of cell cycle (Supplementary Figure 5A). Loss of *Nupr1* altered the genes involved in the regulation of mitotic cell cycle and negative regulation of cell cycle (Supplementary Figure 5B). In addition, the positive regulatory genes of cell cycle, such as *Cdk4, Cdk6, Akt1* and *Akt2*, were upregulated in the *Nupr1*^*-/-*^ HSCs. However, the quiescence regulators of HSCs, such as *Gfi1, Pten, Myb, Hlf, Cdc42* and *Foxo1* were downregulated in the *Nupr1*^*-/-*^ HSCs (Supplementary Figure 5C)^42, 43^. Gene set enrichment analysis (GSEA) illustrated that p53 pathways feedback loops-related genes, including *Trp53, Ccng1, Ctnnb1, Pten*, and *Pik3c2b*, were enriched in WT HSCs (Figure 5A). p53 pathway regulates a series of target genes involving cell cycle arrest, apoptosis, senescence, DNA repair, and metabolism^44^. Interestingly, the expression of p53 was significantly (p < 0.001) reduced to 1/3 of control in *Nupr1*^-/-^ HSCs (Figure 5B). Therefore, we hypothesized that down-regulation of p53 in *Nupr1*^-/-^ HSCs might account for the competitive advantage of the HSCs. MDM2 is a ubiquitin ligase E3 for p53, which is a key repressive regulator of p53 signaling^45^. *Mdm2* deficient mice showed active p53 levels, which is an ideal substitute model of up-regulating p53 since direct overexpressing p53 leading to cell death and blood malignancies in mice^27, 46^. The *Nupr1*^-/-^ mice were crossed to the *Mdm2*^+/-^ mice to achieve up-regulation of p53 expression in *Nupr1*^-/-^ HSCs. The expression level of p53 in *Nupr1*^-/-^ and *Nupr1*^-/-^*Mdm2*^*+/-*^ HSC. The expression level of p53 protein in *Nupr1*^-/-^*Mdm2*^*+/-*^ HSCs is comparable with WT HSCs, which is significantly higher than *Nupr1*^-/-^ HSCs when measured by indirect immunofluorescence assay (Figure 5C, D). In addition, most genes involved in the p53 pathways were up-regulated in the *Nupr1*^*-/-*^*Mdm2*^*+/-*^ HSCs, indicated that the p53 pathway was recovered (Supplementary Figure 6). The Next, we examined the phenotypic HSC of the *Nupr1*^-/-^*Mdm2*^*+/-*^ mice. Flow cytometry analysis showed that *Nupr1*^*-/-*^*Mdm2*^*+/-*^ HSC pool was indistinguishable with *Nupr1*^*-/-*^ and *Mdm2*^*+/-*^ counterparts in terms of ratios and absolute numbers (Figure 6A, B). Further, we tested the competitiveness of *Nupr1*^-/-^*Mdm2*^+/-^ HSCs in parallel with *Nupr1*^-/-^ and *Mdm2*^*+/-*^ HSCs. Two hundred and fifty thousand whole bone marrow nucleated cells from *Nupr1*^-/-^ *Mdm2*^+/-^ mice (CD45.2), *Nupr1*^-/-^ mice (CD45.2) or *Mdm2*^*+/-*^ mice (CD45.2) were transplanted into lethally irradiated recipients (CD45.1) along with equivalent WT (CD45.1) whole bone marrow nucleated cells. In the recipients of *Nupr1*^-/-^*Mdm2*^+/-^ donor cells, the contribution of *Nupr1*^-/-^*Mdm2*^+/-^ cells was significantly (p < 0.001) reduced to ∼20%, which was far below the percentage of *Nupr1*^-/-^ cells in recipients of *Nupr1*^-/-^ donor cells. And the *Mdm2*^*+/-*^ cells only took less than 10% in the PB of recipients 16 weeks after transplantation (Figure 6C). Sixteen weeks after transplantation, we also analyzed the *Nupr1*^-/-^*Mdm2*^+/-^ HSCs in the chimeras. Surprisingly, only a few *Nupr1*^*-/-*^*Mdm2*^*+/-*^ HSCs exists in the HSC pool of the recipients, while the *Nupr1*^*-/-*^ HSCs dominantly occupied in the HSC pool. And the *Mdm2*^+/-^ HSCs almost disappeared in the HSC pool of the recipients (Figure 6D, E). Altogether, the reversion of p53 expression offsets the competitiveness advantage of *Nupr1*^*-/-*^ HSCs.

**Fig 5.**
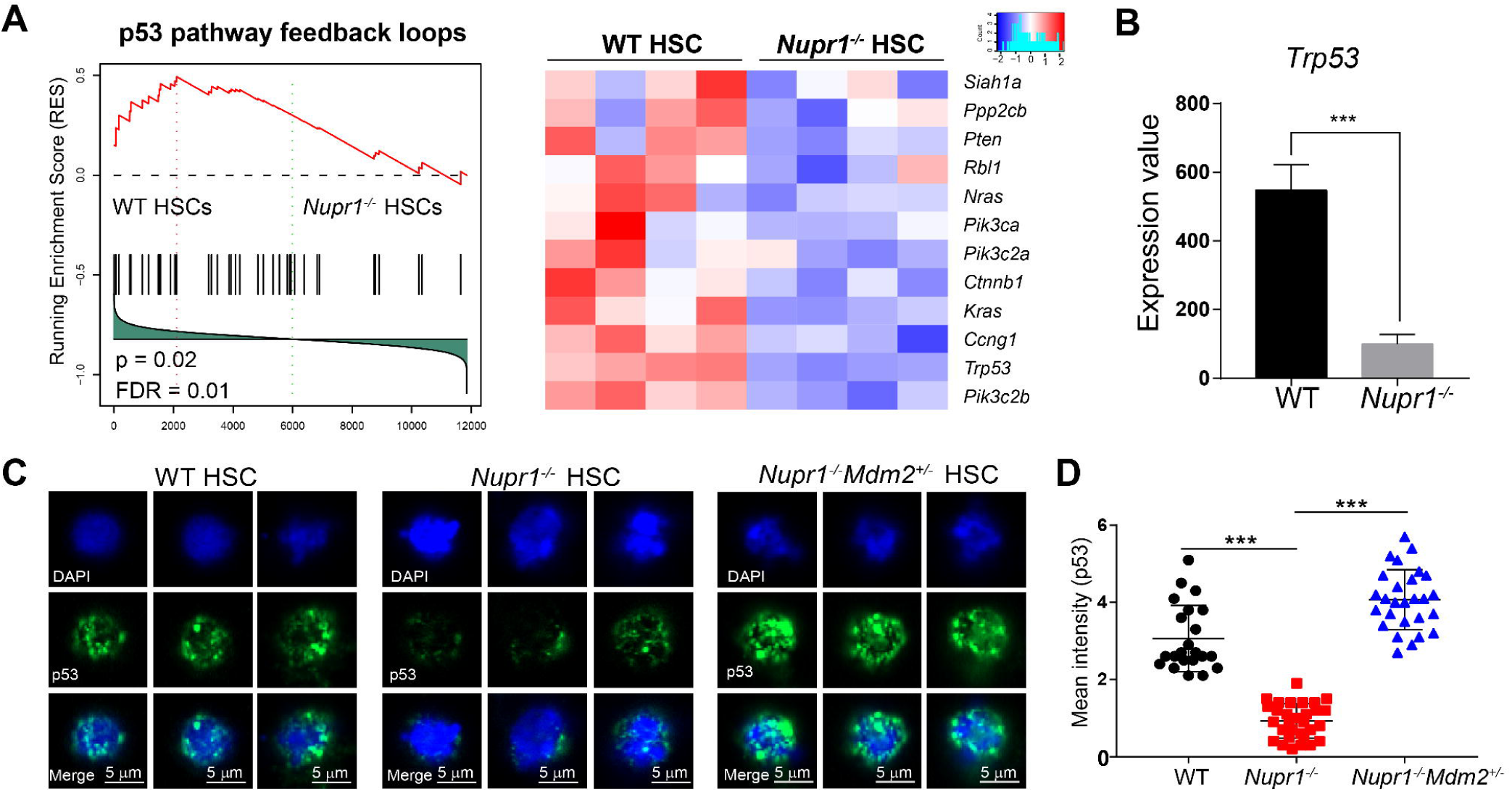
Loss of *Nupr1* confers repopulating advantage on HSCs by regulating p53 check-point signaling. (A) Gene set enrichment analysis (GSEA) of p53 pathway feedback loops in WT HSCs and *Nupr1*^-/-^ HSCs. One thousand HSCs from bone marrow of wild type and *Nupr1*^-/-^ mice were sorted as individual samples for RNA-sequencing. DESeq2 normalized values of the expression data were used for GSEA analysis. Expression of the leading-edge gene subsets was shown. p53 pathway feedback loops-related genes down-regulated in *Nupr1*^-/-^ HSCs (a difference in expression over 1.2-fold; adjusted p value, < 0.05 (DESeq2 R package)). WT HSCs, n = 4 cell sample replicates (one per column); *Nupr1*^-/-^ HSCs, n = 4 cell sample replicates (one per column). (B) Expression level of p*53* in WT HSCs and *Nupr1*^-/-^ HSCs by RNA-seq. Y-axis indicates the expression value (DESeq2 normalized values of the expression data). The expression value (DESeq2 normalized counts) of each gene was illustrated by graphpad. Data are analyzed by unpaired Student’s t-test (two-tailed). ***p < 0.001. Data are represented as mean ± SD (n = 4 mice for each group). (C) Immunofluorescence measurement of p53 proteins in single HSCs from the WT, *Nupr1*^*-/-*^, *Mdm2*^*+/-*^*Nupr1*^*-/-*^ mice. Images of three representative single cell of each group are shown. (D) Mean intensity of p53 fluorescence in WT, *Nupr1*^*-/-*^, *Mdm2*^*+/-*^*Nupr1*^*-/-*^ HSCs. Each dot represents a single cell. Data are analyzed by One-way ANOVA. *\*\*\**p<0.001. WT, n=22; *Nupr1*^*-/-*^, n=30; *Mdm2*^*+/-*^*Nupr1*^*-/-*^, n=27. Data are represented as mean ± SD.

**Fig 6.**
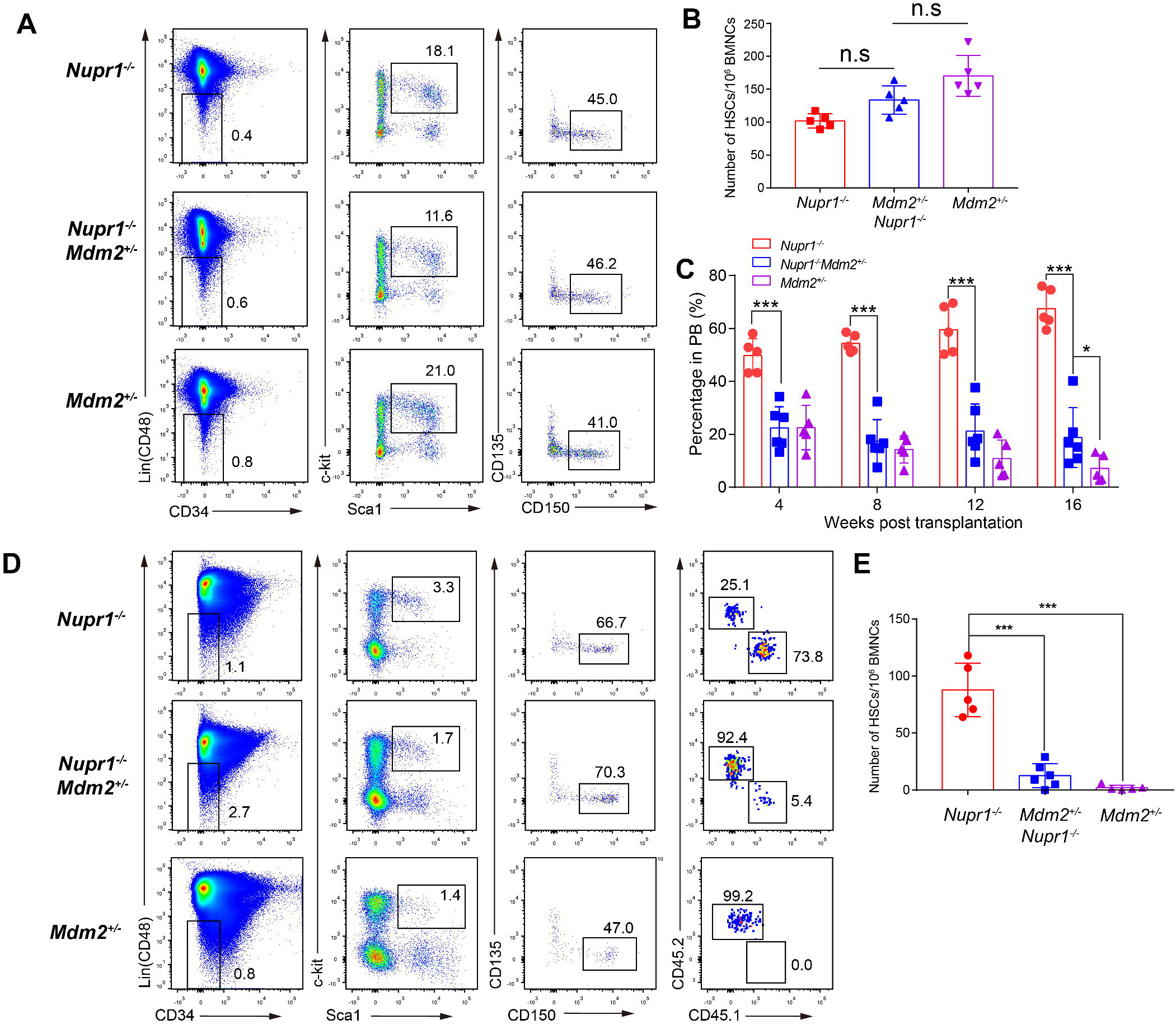
Reversion of p53 expression by allelic depletion of Mdm2 gene offsets the repopulating advantage of *Nupr1*^-/-^ HSCs. (A) Representative plots of HSC analysis by flow cytometry from *Nupr1*^*-/-*^, *Nupr1*^*-/-*^*Mdm2*^*+/-*^ and *Mdm2*^*+/-*^ mice bone marrow. (B) Statistical analysis of *Nupr1*^*-/-*^, *Nupr1*^*-/-*^*Mdm2*^*+/-*^ and *Mdm2*^*+/-*^ HSC number. Data are analyzed by One-way ANOVA. n=5. (C) Donor bone marrow cells (2.5×10^5^) from *Nupr1*^-/-^ (red), *Nupr1*^-/-^*Mdm2*^+/-^ (blue) (CD45.2) or *Mdm2*^+/-^ (purple) mice were transplanted into lethally irradiated recipient mice (CD45.1) along with 2.5×10^5^ recipient bone marrow cells. Data are analyzed by unpaired Student’s t-test. *p < 0.05, ***p < 0.001. Data are represented as mean ± SD. *Nupr1*^-/-^, n = 5 mice; *Nupr1*^-/-^*Mdm2*^+/-^, n = 6 mice; Mdm2^+/-^: n=5. (D) Flow cytometry analysis of donor-derived HSCs and recipient HSCs in bone marrow of recipient mice at four months after transplantation. HSCs were gated as Lin (CD2, CD3, CD4, CD8, B220, Gr1, CD11b, Ter119^-^) CD48^-^ Sca1^+^ c-Kit^+^ CD150^+^ CD34^-^ CD135^-^. Plots from one representative mice of each group are shown. (E) Statistical analysis of donor-derived HSC percentage in recipient mice at four months after transplantation. Data are analyzed by One-way ANOVA. ***p < 0.001. Data are represented as mean ± SD. *Nupr1*^-/-^, n = 5 mice; *Nupr1*^-/-^*Mdm2*^+/-^, n = 6 mice; Mdm2^+/-^: n=5.

## Discussion

The intrinsic networks of regulating the quiescence of HSCs are largely unknown. In this study, loss of *Nupr1* (p8), a gene preferentially expressed in long-term HSCs, tunes the quiescence threshold of HSCs on homeostasis condition without compromising their key functions in hematopoiesis. *Nupr1* coordinates with p53 to form a signaling machinery regulating HSC quiescence and turnover rate. For the first time, we unveil the *de novo* role of *Nupr1* in controlling HSC quiescence.

*Nupr1*^-/-^ HSCs replenished faster than WT HSCs under homeostasis. However, the size of *Nupr1*^*-/-*^ HSC pool was not altered. These data implicate that despite the existence of intrinsic machinery of controlling HSC quiescence, the scale of HSC-pool is restricted by extrinsic bone marrow microenvironment^47^. Conventionally, molecules activating HSCs showed transiently phenotypic proliferation of HSCs but eventually led to their functional exhaustion and even tumors^36-40^. Interestingly, *Nupr1* signaling seemingly plays a unique role in regulating HSC quiescence and turnover rates, as deletion of *Nupr1* maintains the hematopoiesis features of HSCs. Consistently, enforced CDK6 expression in HSCs confers competitive advantage without impairing their stemness and multi-lineage potential^9^. These evidence supports the concept that targeting the intrinsic machinery of balancing HSC quiescence threshold might safely promoting engraftment.

Loss of *Nupr1* in HSCs resulted in engraftment advantage. Under the transplantation stress settings, the HSC niche occupied by WT HSCs was ablated, providing niche vacuum for donor *Nupr1*^*-/-*^ HSC entrance. The dominance of *Nupr1*^-/-^ HSCs is a con-sequence of fast turnover rate of these cells over WT counterparts. In the previous re-search, loss of Dnmt3a also leads to clonal dominance of HSCs, however, accompanied with hematopoiesis failure due to differentiation block^4, 48^. Thus, the engraftment advantage caused by loss of *Nupr1* might have prospective translational implications for hematopoietic stem cells transplantation (HSCT), since a faster recovery of hema-topoiesis in transplanted host definitely reduces infection risks in patients^49, 50^.

In our models, *Nupr1* regulated hematopoietic homeostasis via targeting p53 pathway. Consistently, p53 is essential in regulating hematopoietic homeostasis^27^. Whether NUPR1 directly interacts with p53 in HSC context remain unknown, as currently antibodies suitable for protein-protein interaction assays are not available. NUPR1 and p53 directly interacted in human breast epithelial cells^22^. Knocking out p53 in HSCs can promote HSC expansion, but directly targeting p53 caused HSC apoptosis and tumorigenesis^51^. Thus, *Nupr1* might behave as an upstream regulator of p53 signaling and uniquely regulate cell quiescence in HSC context. In previous research, Mdm2 is a key repressive regulator of p53 signaling. The complete deletion of Mdm2 will lead to embryo-lethal because of the excess expression of p53^46^. Therefore, we cross the *Nupr1*^*-/-*^ mice with the *Mdm2*^*+/-*^ to up-regulate the p53 expression.

In Vav-Cre crossed mice model, *Nupr1* was deleted from embryonic stage. To exclude the possibility that the effect seen in the adulthood is a consequence of an effect coming from the embryo, mice bearing the *Nupr1* floxp-floxp insertion was crossed with Mx1-Cre mice to generate the induced knockout mice of *Nupr1*. The deletion of *Nupr1* gene in the Nupr1^fl/fl^Mx1-cre mice was verified by PCR result of the genome DNA from the BM cells and the qPCR result of Nupr1 expression level from the HSCs (Supplementary Figure 7A, B). Cre expression was induced through intraperitoneal injection of polyinosinic-polycytidylic acid (250μg/mice) every other day one week before analysis. More Nupr1^fl/fl^Mx1-cre HSCs entered into G1-S-S2 and M phase (median value: 26.35%) than the littermates Control mice (median value: 14.13%) (Supplementary Figure 7C-E). The competitive transplantation result showed that Nupr1^fl/fl^Mx1-cre cells took advantage in the PB of recipients (60%-80%), while control cells maintained 50%-60% in the chimeras (Supplementary Figure 7F). To further explore whether Nupr1^fl/fl^Mx1-cre HSCs dominate in recipients, we sacrificed the chimeras and analyzed the HSCs 16 weeks after transplantation. Compared with the control HSCs, the proportion and absolute number of Nupr1^fl/fl^Mx1-cre HSCs were significantly more (∼2 folds) than the control HSC competitors in primary recipients (Supplementary figure 7G, H). Taken together, the deletion of *Nupr1* in the adulthood is not a consequence of an effect coming from the embryo.

In conclusion, loss of *Nupr1* in HSCs promotes engraftment by tuning the quiescence threshold of HSCs via regulating p53 check-point pathway. Our study unveils the prospect of shortening the engraftment time of HSCT by targeting the intrinsic machinery of controlling HSC quiescence.

## Acknowledgments

This work was supported by grants from the National Natural Science Foundation of China (31900814, 81925002, 81922002), Strategic Priority Research Program of the Chinese Academy of Sciences (XDA16010601), Key Research & Development Program of Guangzhou Regenerative Medicine and Health Guangdong Laboratory (2018GZR110104006), CAS Key Research Program of Frontier Sciences (QYZDB-SSW-SM057), Science and Technology Planning Project of Guangdong Province (2017B030314056).

## Author contributions

T.J.W. and C.X.X. performed research, analyzed data and wrote the paper; Y.D. and Q.T.W. analyzed RNA-Seq data; S.H., F.D., K.T.W., X.F.L., L.J.L., Y.G., and Y.X.G. performed experiments; J.D., T.C., and H.C. discussed the manuscript; J.Y.W. designed research, and wrote the manuscript.

## References

1. Cheshier SH, Morrison SJ, Liao X, Weissman IL. In vivo proliferation and cell cycle kinetics of long-term self-renewing hematopoietic stem cells. Proc Natl Acad Sci U S A. 1999;96(6):3120–3125.

2. Wilson A, Laurenti E, Oser G, et al. Hematopoietic stem cells reversibly switch from dormancy to self-renewal during homeostasis and repair. Cell. 2008;135(6):1118–1129.

3. Rodrigues NP, Janzen V, Forkert R, et al. Haploinsufficiency of GATA-2 perturbs adult hematopoietic stem-cell homeostasis. Blood. 2005;106(2):477–484.

4. Challen GA, Sun D, Jeong M, et al. Dnmt3a is essential for hematopoietic stem cell differentiation. Nat Genet. 2011;44(1):23–31.

5. Mayle A, Yang L, Rodriguez B, et al. Dnmt3a loss predisposes murine hematopoietic stem cells to malignant transformation. Blood. 2015;125(4):629–638.

6. Santaguida M, Schepers K, King B, et al. JunB protects against myeloid malignancies by limiting hematopoietic stem cell proliferation and differentiation without affecting self-renewal. Cancer Cell. 2009;15(4):341–352.

7. Takubo K, Goda N, Yamada W, et al. Regulation of the HIF-1alpha level is essential for hematopoietic stem cells. Cell Stem Cell. 2010;7(3):391–402.

8. Tesio M, Tang Y, Mudder K, et al. Hematopoietic stem cell quiescence and function are controlled by the CYLD-TRAF2-p38MAPK pathway. J Exp Med. 2015;212(4):525–538.

9. Laurenti E, Frelin C, Xie S, et al. CDK6 levels regulate quiescence exit in human hematopoietic stem cells. Cell Stem Cell. 2015;16(3):302–313.

10. Mallo GV, Fiedler F, Calvo EL, et al. Cloning and expression of the rat p8 cDNA, a new gene activated in pancreas during the acute phase of pancreatitis, pancreatic development, and regeneration, and which promotes cellular growth. J Biol Chem. 1997;272(51):32360–32369.

11. Ree AH, Tvermyr M, Engebraaten O, et al. Expression of a novel factor in human breast cancer cells with metastatic potential. Cancer Res. 1999;59(18):4675–4680.

12. Ree AH, Pacheco MM, Tvermyr M, Fodstad O, Brentani MM. Expression of a novel factor, com1, in early tumor progression of breast cancer. Clin Cancer Res. 2000;6(5):1778–1783.

13. Ito Y, Yoshida H, Motoo Y, et al. Expression and cellular localization of p8 protein in thyroid neoplasms. Cancer Lett. 2003;201(2):237–244.

14. Mohammad HP, Seachrist DD, Quirk CC, Nilson JH. Reexpression of p8 contributes to tumorigenic properties of pituitary cells and appears in a subset of prolactinomas in transgenic mice that hypersecrete luteinizing hormone. Mol Endocrinol. 2004;18(10):2583–2593.

15. Brannon KM, Million Passe CM, White CR, Bade NA, King MW, Quirk CC. Expression of the high mobility group A family member p8 is essential to maintaining tumorigenic potential by promoting cell cycle dysregulation in LbetaT2 cells. Cancer Lett. 2007;254(1):146–155.

16. Jiang WG, Davies G, Martin TA, Kynaston H, Mason MD, Fodstad O. Com-1/p8 acts as a putative tumour suppressor in prostate cancer. Int J Mol Med. 2006;18(5):981–986.

17. Malicet C, Lesavre N, Vasseur S, Iovanna JL. p8 inhibits the growth of human pancreatic cancer cells and its expression is induced through pathways involved in growth inhibition and repressed by factors promoting cell growth. Mol Cancer. 2003;2(37.

18. Malicet C, Giroux V, Vasseur S, Dagorn JC, Neira JL, Iovanna JL. Regulation of apoptosis by the p8/prothymosin alpha complex. Proc Natl Acad Sci U S A. 2006;103(8):2671–2676.

19. Vasseur S, Hoffmeister A, Garcia-Montero A, et al. p8-deficient fibroblasts grow more rapidly and are more resistant to adriamycin-induced apoptosis. Oncogene. 2002;21(11):1685–1694.

20. Carracedo A, Lorente M, Egia A, et al. The stress-regulated protein p8 mediates cannabinoid-induced apoptosis of tumor cells. Cancer Cell. 2006;9(4):301–312.

21. Gironella M, Malicet C, Cano C, et al. p8/nupr1 regulates DNA-repair activity after double-strand gamma irradiation-induced DNA damage. J Cell Physiol. 2009;221(3):594–602.

22. Clark DW, Mitra A, Fillmore RA, et al. NUPR1 interacts with p53, transcriptionally regulates p21 and rescues breast epithelial cells from doxorubicin-induced genotoxic stress. Curr Cancer Drug Targets. 2008;8(5):421–430.

23. Dumble M, Moore L, Chambers SM, et al. The impact of altered p53 dosage on hematopoietic stem cell dynamics during aging. Blood. 2007;109(4):1736–1742.

24. Lotem J, Sachs L. Hematopoietic cells from mice deficient in wild-type p53 are more resistant to induction of apoptosis by some agents. Blood. 1993;82(4):1092–1096.

25. Shounan Y, Dolnikov A, MacKenzie KL, Miller M, Chan YY, Symonds G. Retroviral transduction of hematopoietic progenitor cells with mutant p53 promotes survival and proliferation, modifies differentiation potential and inhibits apoptosis. Leukemia. 1996;10(10):1619–1628.

26. Bondar T, Medzhitov R. p53-mediated hematopoietic stem and progenitor cell competition. Cell Stem Cell. 2010;6(4):309–322.

27. Liu Y, Elf SE, Miyata Y, et al. p53 regulates hematopoietic stem cell quiescence. Cell Stem Cell. 2009;4(1):37–48.

28. Chen J, Ellison FM, Keyvanfar K, et al. Enrichment of hematopoietic stem cells with SLAM and LSK markers for the detection of hematopoietic stem cell function in normal and Trp53 null mice. Exp Hematol. 2008;36(10):1236–1243.

29. Wang YV, Leblanc M, Fox N, et al. Fine-tuning p53 activity through C-terminal modification significantly contributes to HSC homeostasis and mouse radiosensitivity. Genes Dev. 2011;25(13):1426–1438.

30. Liu D, Ou L, Clemenson GD, Jr., et al. Puma is required for p53-induced depletion of adult stem cells. Nat Cell Biol. 2010;12(10):993–998.

31. Yamashita M, Nitta E, Suda T. Regulation of hematopoietic stem cell integrity through p53 and its related factors. Ann N Y Acad Sci. 2016;1370(1):45–54.

32. Wilkinson AC, Ishida R, Kikuchi M, et al. Long-term ex vivo haematopoietic-stem-cell expansion allows nonconditioned transplantation. Nature. 2019;571(7763):117–121.

33. Yamamoto R, Morita Y, Ooehara J, et al. Clonal analysis unveils self-renewing lineage-restricted progenitors generated directly from hematopoietic stem cells. Cell. 2013;154(5):1112–1126.

34. Hu Y, Smyth GK. ELDA: extreme limiting dilution analysis for comparing depleted and enriched populations in stem cell and other assays. J Immunol Methods. 2009;347(1-2):70–78.

35. Kiel MJ, He S, Ashkenazi R, et al. Haematopoietic stem cells do not asymmetrically segregate chromosomes or retain BrdU. Nature. 2007;449(7159):238–242.

36. Motoda L, Osato M, Yamashita N, et al. Runx1 protects hematopoietic stem/progenitor cells from oncogenic insult. Stem Cells. 2007;25(12):2976–2986.

37. Miyamoto K, Araki KY, Naka K, et al. Foxo3a is essential for maintenance of the hematopoietic stem cell pool. Cell Stem Cell. 2007;1(1):101–112.

38. Ficara F, Murphy MJ, Lin M, Cleary ML. Pbx1 regulates self-renewal of long-term hematopoietic stem cells by maintaining their quiescence. Cell Stem Cell. 2008;2(5):484–496.

39. Tipping AJ, Pina C, Castor A, et al. High GATA-2 expression inhibits human hematopoietic stem and progenitor cell function by effects on cell cycle. Blood. 2009;113(12):2661–2672.

40. Campbell TB, Basu S, Hangoc G, Tao W, Broxmeyer HE. Overexpression of Rheb2 enhances. mouse hematopoietic progenitor cell growth while impairing stem cell repopulation. Blood. 2009;114(16):3392–3401.

41. Stobbe CC, Park SJ, Chapman JD. The radiation hypersensitivity of cells at mitosis. Int J Radiat Biol. 2002;78(12):1149–1157.

42. Hao S, Chen C, Cheng T. Cell cycle regulation of hematopoietic stem or progenitor cells. Int J Hematol. 2016;103(5):487–497.

43. Pietras EM, Warr MR, Passegue E. Cell cycle regulation in hematopoietic stem cells. J Cell Biol. 2011;195(5):709–720.

44. Li T, Kon N, Jiang L, et al. Tumor suppression in the absence of p53-mediated cell-cycle arrest, apoptosis, and senescence. Cell. 2012;149(6):1269–1283.

45. Honda R, Tanaka H, Yasuda H. Oncoprotein MDM2 is a ubiquitin ligase E3 for tumor suppressor p53. FEBS Lett. 1997;420(1):25–27.

46. Abbas HA, Maccio DR, Coskun S, et al. Mdm2 is required for survival of hematopoietic stem cells/progenitors via dampening of ROS-induced p53 activity. Cell Stem Cell. 2010;7(5):606–617.

47. Anthony BA, Link DC. Regulation of hematopoietic stem cells by bone marrow stromal cells. Trends Immunol. 2014;35(1):32–37.

48. Challen GA, Sun D, Mayle A, et al. Dnmt3a and Dnmt3b have overlapping and distinct functions in hematopoietic stem cells. Cell Stem Cell. 2014;15(3):350–364.

49. Young JH, Logan BR, Wu J, et al. Infections after Transplantation of Bone Marrow or Peripheral Blood Stem Cells from Unrelated Donors. Biol Blood Marrow Transplant. 2016;22(2):359–370.

50. Safdar A, Armstrong D. Infections in patients with hematologic neoplasms and hematopoietic stem cell transplantation: neutropenia, humoral, and splenic defects. Clin Infect Dis. 2011;53(8):798–806.

51. Orazi A, Kahsai M, John K, Neiman RS. p53 overexpression in myeloid leukemic disorders is associated with increased apoptosis of hematopoietic marrow cells and ineffective hematopoiesis. Mod Pathol. 1996;9(1):48–52.

