## Supplementary Methods and figures for "Loss of *Nupr1* promotes engraftment by tuning the quiescence threshold of hematopoietic stem cell repository via regulating p53-checkpoint pathway"

**Supplementary information**

### Supplementary Methods

#### Flow cytometry analysis

For HSC staining, total BM cells were stained with antibodies against (CD2, CD3, CD4, CD8, Ter119, B220, Gr1, CD48)-Alexa Flour700, Sca1-Percp-cy5.5, c-kit-APC-cy7, CD150-PE-cy7, CD34-FITC and CD135-PE[1-4]. Cells were analyzed by LSR Fortessa (BD Bioscience). For lineage analysis of peripheral blood, the white blood cells were stained with antibodies of anti-CD45.1-FITC, anti-CD45.2-percp-cy5.5, anti-CD90.2-APC, anti-CD19-PE, anti-CD11b-PE-cy7, anti-Gr-1-APC-eFlour® 780.

**RNA-Seq and data analysis.** For HSC library preparation, HSCs ($Lin^-CD48^-Sca1^+cKit^+CD150^+$) were sorted from 8-10 weeks old *Nupr1*$^{-/-}$ mice and wild type mice. HSCs were sorted from four mice of each group. 1000 target cells per sample were sorted into 500 µl DPBS-BSA buffer (0.5%BSA) using 1.5ml EP tube and transferred into 250 µl tube to spin down with 500 g. The cDNA of sorted 1000-cell aliquots were generated and amplified as described previously[5]. The qualities of the amplified cDNA were examined by Q-PCR analysis of housekeeping genes (*B2m*, *Actb*, *Gapdh*, *Ecf1a1*). Samples passed quality control were used for sequencing library preparation by illumina Nextera XT DNA Sample Preparation Kit (FC-131-1096).

For data analysis, all libraries were sequenced by illumina sequencers NextSeq 500. The fastq files of sequencing raw data samples were generated using illumina bcl2fastq software (version: 2.16.0.10) and were uploaded to Gene Expression Omnibus public database (GSE131071). Raw reads were aligned to mouse genome (mm10) by HISAT2[6] (version: 2.1.0) as reported. And raw counts were calculated by featureCounts of subread[7] (version 1.6.0). Differential gene expression analysis was performed by DESeq2[8] (R package version: 1.18.1). Heatmaps were plotted using gplots (R package, version 3.01). GSEA was performed as described[9]. The gene set (p53 pathway feedback loop) for GSEA were from PANTHER pathways dataset.

**Quantitative real-time PCR**

Total RNA was extracted from ten thousand purified HSCs and MPPs with an RNeasy micro kit (QIAGEN). Then, 2 ng of RNA was used for linear amplification according to the manufacturer's instructions (3302–12, Ovation Pico WTA System V2, NuGEN Technologies, Inc.). The RNA was diluted and 10ng RNA was used as the template for quantitative real-time PCR (CFX-96, Bio-Rad). The forward primer of *Nupr1* is 5'-CCCTTCCCAGCAACCTCTAA-3'and the reverse primer is 5'-AGCTTCTCTCTTGGTCCGAC-3'. Fold expression relative to the reference gene was calculated using the comparative method $2^{-\Delta\Delta Ct}$, and the values were normalized to 1 for comparison.

**Indirect Immunofluorescence Assay**

Sorted HSCs were directly pipetted onto the poly-lysine coated slides (100-500 cells in 5μl) and incubated at room temperature for 10 min. Upon the solution was completely dry, the cells were fixed with 4%PFA for 10 min following with 0.15%Triton X-100 permeabilization for 2 min at room temperature. To avoid non-specific antibody binding, the cells were blocked in 1% BSA/PBS for 1-2h at room temperature and then incubated with the primary p53 antibody in 1% BSA in PBS overnight at $4^{o}C$ (Abcam, ab16465). Slides were washed three times in PBS and incubated with secondary antibodies for 1h at room temperature in 1% BSA in PBS (donkey anti-mouse Alexa Fluor® 488, Abcam, ab150105). After washing the slides, the cells were incubated with DAPI solution for 10 min. Confocal analysis was performed at high resolution with a Zeiss laser scanning confocal microscope, LSM-800. The images were processed with ZEN 2012 software (blue edition).

### Statistical analysis

Normal distribution of data was tested with SPSS applying Shapiro-Wilk normality test (SPSS v.23, IBM Corp., Armonk, NY, USA). The data were represented as mean ± SD. Two-tailed independent or dependent Student's t-tests were performed for comparison of two groups of data. For the analysis of three groups or more, one-way ANOVA was used, and further significance analysis among groups was analyzed by Post Hoc Test (equal variances, Turkey-HSD; unequal variances, Games-Howell). P values of less than 0.05 were considered statistically significant (*$p < 0.05$, **$p < 0.01$, ***$p < 0.001$).

### Supplementary Figures

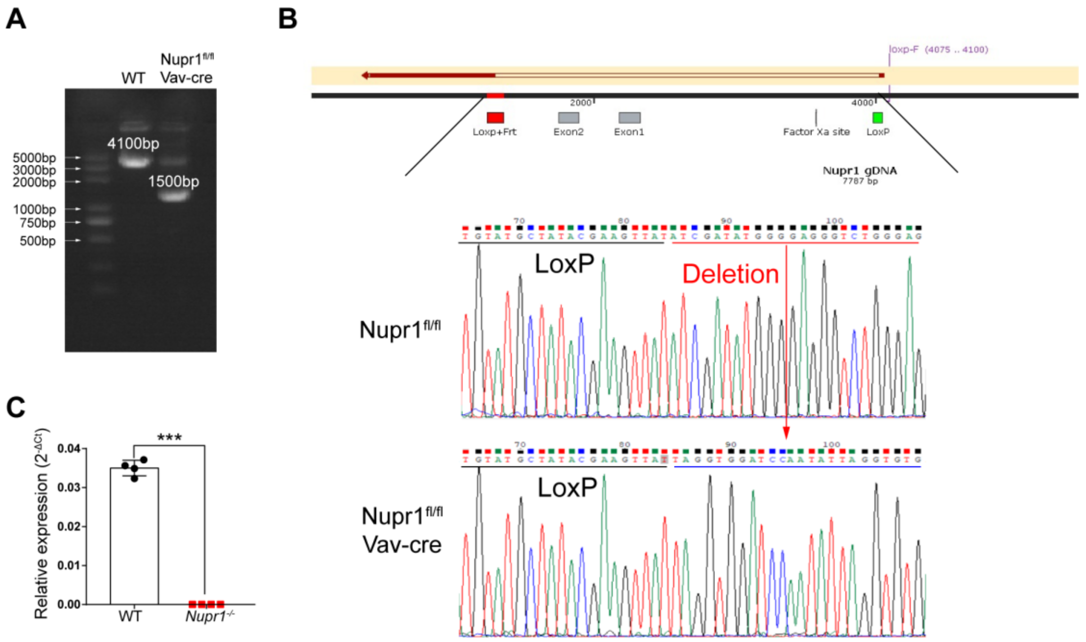


**Supplementary fig 1. Validation of Nupr1 knockout in the Nupr1$^{fl/fl}$Vav-cre mice**

(A) PCR verification of Nupr1 gene from the Nupr1$^{fl/fl}$Vav-cre mice. The genome DNA of BM cells were extracted from WT and Nupr1$^{fl/fl}$Vav-cre mice. And the Nupr1 gene was amplified by specific primers. Normal Nupr1 gene length: 4100bp, Nupr1 deleted fragment: 1500bp.

(B) Sequencing of *Nupr1* gene which was amplified from Nupr1$^{fl/fl}$ BM and Nupr1$^{fl/fl}$Vav-cre BM.

(C) Real-time qPCR validation of Nupr1 knockout in the Nupr1$^{fl/fl}$Vav-cre HSCs. RNA from 500-1000 sorted WT or Nupr1$^{fl/fl}$Vav-cre HSCs were extracted and reverse transcript into cDNA. ***p<0.001, n=4

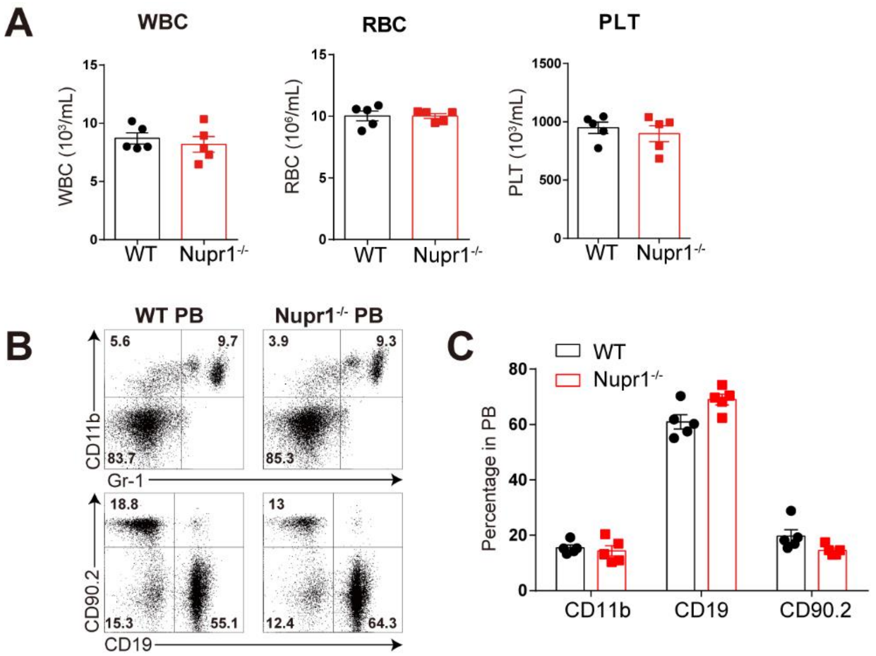


**Supplementary fig 2. The hematopoiesis of *Nupr1*$^{-/-}$ mice was normal**

(A) Complete blood counts of peripheral blood from WT mice and *Nupr1*$^{-/-}$ mice. Eight-ten-week-old mice were used for detection. WBC: white blood cells; RBC: red blood cells; PLT: platelets.

(B) Flow cytometry analysis of lineage cells in peripheral blood from WT mice and *Nupr1*$^{-/-}$ mice. Myeloid cells were detected by CD11b$^{+}$; Granulocytes were detected by Gr-1$^{+}$; B lymphocytes were detected by CD19$^{+}$; T lymphocytes were detected by CD90.2$^{+}$.

(C) Statistical analysis of lineage percentage in peripheral blood from WT mice and *Nupr1*$^{-/-}$ mice.

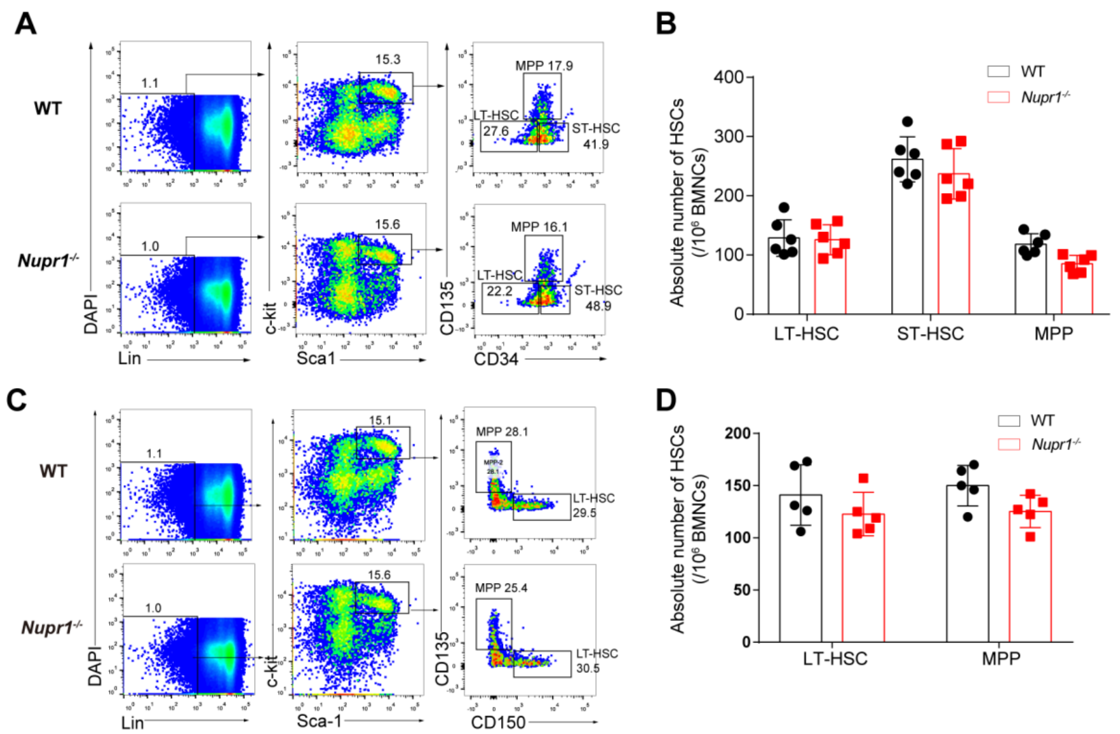


**Supplementary fig 3. The number of LT-HSCs were compatible between WT and *Nupr1*$^{-/-}$ HSC**

(A) Representative plots of LT-HSCs, ST-HSCs and MPPs in the *Nupr1*$^{-/-}$ mice by flow cytometry analysis. Long-term HSCs (LT-HSCs) were defined as Lin$^{-}$Sca-1$^{+}$c-Kit$^{+}$CD135$^{-}$CD34$^{-}$, progenitors including short-term HSCs (ST-HSCs) as Lin$^{-}$Sca-1$^{+}$c-Kit$^{+}$CD135$^{-}$CD34$^{+}$, and multipotent progenitors (MPPs) as Lin$^{-}$Sca-1$^{+}$c-Kit$^{+}$CD135$^{high}$CD34$^{+}$.

(B) Statistic analysis of LT-HSCs, ST-HSCs and MPPs in the *Nupr1*$^{-/-}$ mice using the combination marker of LSK, CD135 and CD34.

(C) Representative plots of LT-HSCs and MPPs in the *Nupr1*$^{-/-}$ mice by flow cytometry analysis. Long-term HSCs (LT-HSCs) were defined as Lin$^{-}$Sca-1$^{+}$c-Kit$^{+}$CD135$^{-}$CD150$^{+}$ and multipotent progenitors (MPPs) as Lin$^{-}$Sca-1$^{+}$c-Kit$^{+}$CD135$^{high}$CD150$^{-}$.

(D) Statistic analysis of LT-HSCs and MPPs in the *Nupr1*$^{-/-}$ mice using the combination marker of LSK, CD135 and CD150.

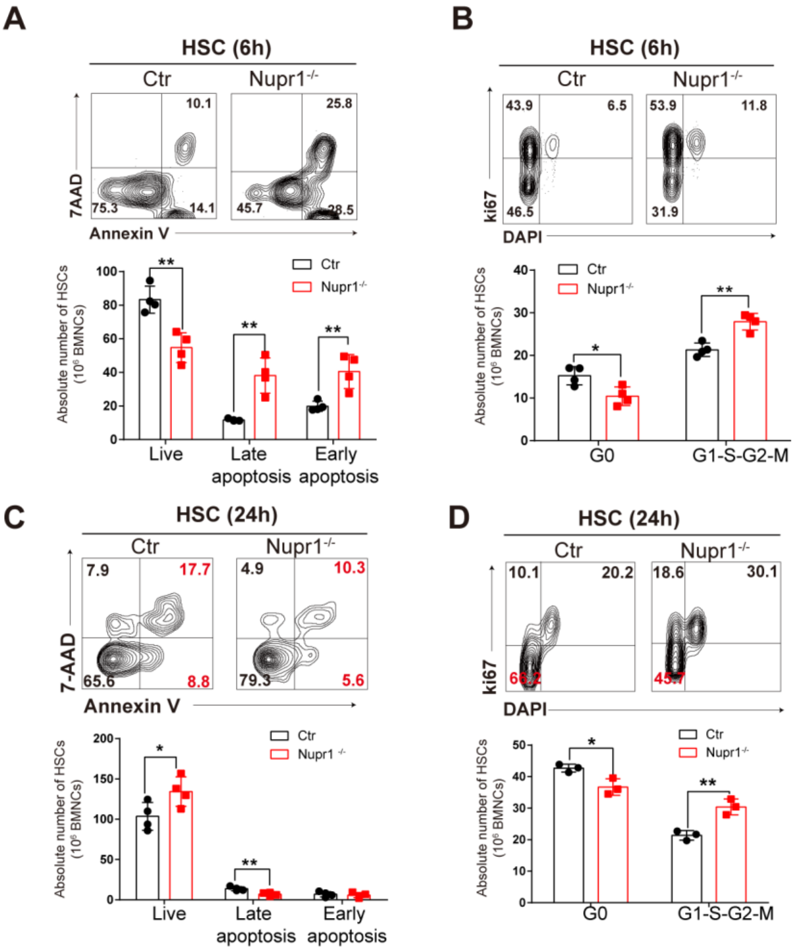


**Supplementary fig 4. Nupr1$^{-/-}$ HSC are susceptible to irradiation-stress but recover fast**

Eight-week-old WT mice and Nupr1$^{-/-}$ mice were irradiated by a single dose of 4Gy irradiation (rate = 1Gy/min). Apoptosis and cell cycle of the irradiated mice were analyzed after 6h and 24h respectively.

(A) Apoptotic analysis of Nupr1$^{-/-}$ and WT HSCs 6 hours after irradiation by flow cytometry. Apoptotic plots of HSCs from representative WT and Nupr1 KO mice were shown. Live cells: AnnexinV-7AAD-; Early apoptosis cells: AnnexinV+7AAD-; Late apoptosis cells: AnnexinV+7AAD+. *$p < 0.05$, ***$p < 0.001$. Unpaired Student's t-test (two-tailed). Data are represented as mean ± SD (n = 4 mice for each group).

(B) Cell cycle analysis of Nupr1$^{-/-}$ and WT HSCs 6 hours after irradiation by flow cytometry. Cell cycle plots of HSCs from representative WT and Nupr1$^{-/-}$ mice were shown. G0 (Ki-67$^{low}$DAPI$^{2N}$), G1 (Ki-67$^{high}$DAPI$^{2N}$), G2-S-M (Ki-67$^{high}$DAPI$^{>2N-4N}$). *$p < 0.05$, **$p < 0.01$. Unpaired Student's t-test (two-tailed). Data are represented as mean ± SD (n = 4 mice for each group).

(C) Apoptotic analysis of Nupr1$^{-/-}$ and WT HSCs 24 hours after irradiation by flow cytometry. Apoptotic plots of HSCs from representative WT and Nupr1$^{-/-}$ mice were shown. Live cells: AnnexinV-7AAD-; Early apoptosis cells: AnnexinV+7AAD-; Late apoptosis cells: AnnexinV+7AAD+. *$p < 0.05$, **$p < 0.01$. Unpaired Student's t-test (two-tailed). Data are represented as mean ± SD (n = 4 mice for each group).

(D) Cell cycle analysis of Nupr1$^{-/-}$ and WT HSCs 24 hours after irradiation by flow cytometry. Cell cycle plots of HSCs from representative WT and Nupr1$^{-/-}$ mice were shown. G0 (Ki-67$^{low}$DAPI$^{2N}$), G1 (Ki-67$^{high}$DAPI$^{2N}$), G2-S-M (Ki-67$^{high}$DAPI$^{>2N-4N}$). ***$p < 0.001$. Unpaired Student's t-test (two-tailed). Data are represented as mean ± SD (n = 4 mice for each group).

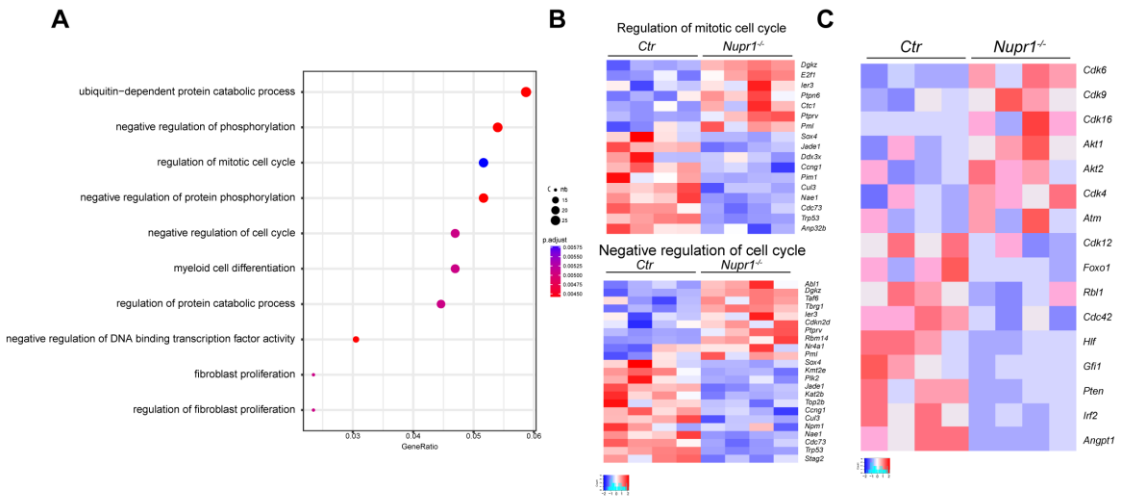


**Supplementary fig 5. RNAseq analysis of Nupr1$^{-/-}$ HSCs**

(A) Gene ontology (GO)–enrichment analysis of the two-fold differentially expressed genes. Each symbol represents a GO term (noted in plot); color indicates adjusted P value (Padj (significance of the GO term); right), and symbol size is proportional to the number of genes.

(B) Expression of genes involved in the regulation of mitotic cell cycle and negative regulation of cell cycle which selected from the list of differentially expressed genes (a difference in expression of over 2-fold; adjusted P value < 0.05 (DESeq2 R package)). n=4.

(C) Expression of selected genes involved in regulating HSC cell cycle and quiescence. One biological replicate per column.

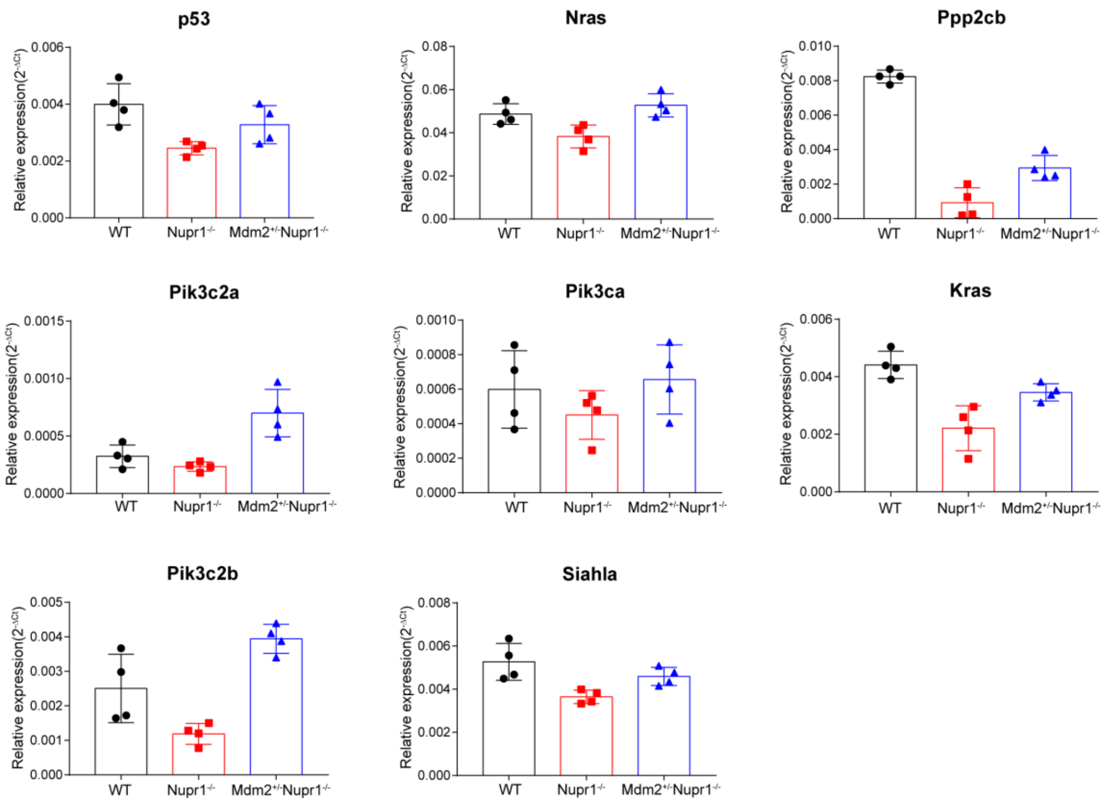


**Supplementary fig 6. The genes involved in the p53 related pathways were recovered in the *Nupr1$^{-/-}$Mdm2$^{+/-}$* mice**

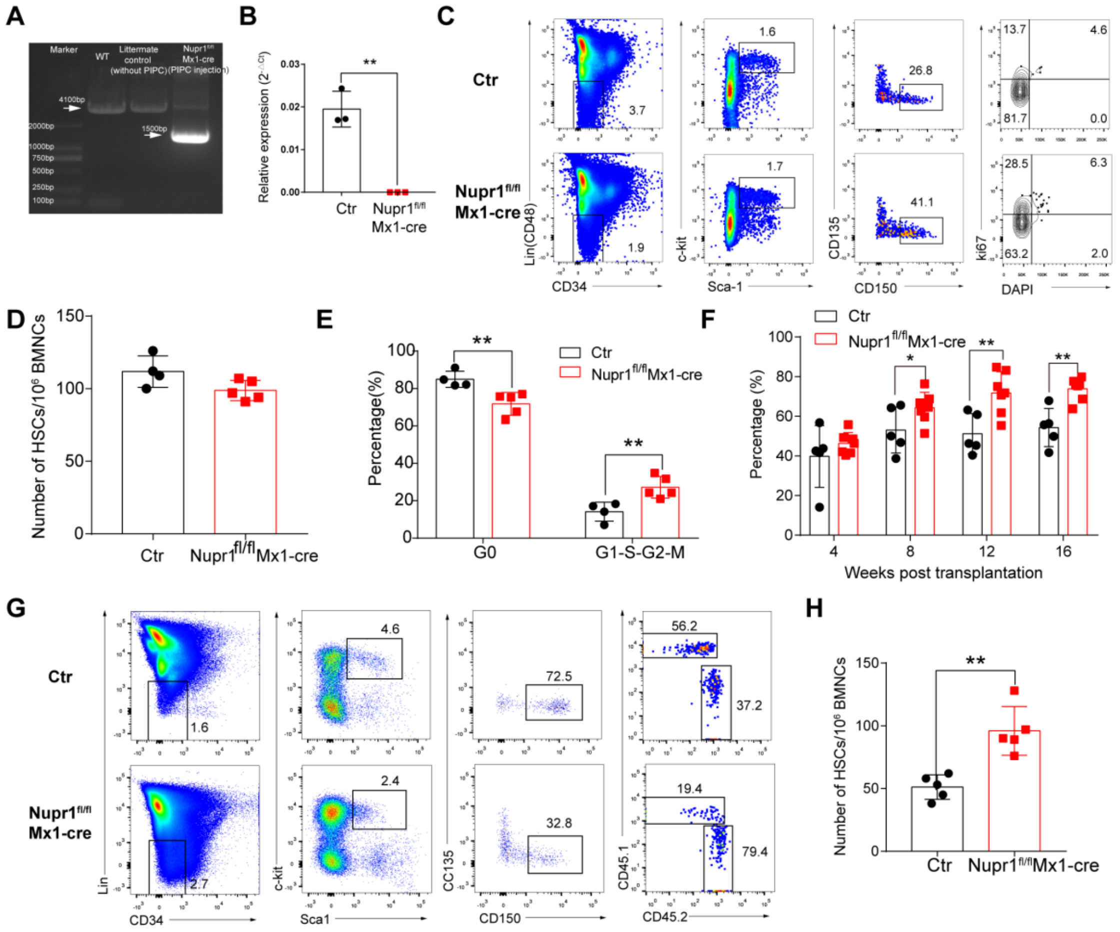


**Supplementary fig 7. The deletion of Nupr1 in the adulthood also promotes engraftment**

(A) PCR verification of Nupr1 gene from the Nupr1$^{fl/fl}$Mx1-cre mice. The genome DNA of BM cells were extracted from WT and Nupr1$^{fl/fl}$Mx1-cre mice. And the Nupr1 gene was amplified by specific primers. Normal Nupr1 gene length: 4100bp, Nupr1 deleted fragment: 1500bp.

(B) Real-time qPCR validation of Nupr1 knockout in the Nupr1$^{fl/fl}$Mx1-cre HSCs. RNA from 500-1000 sorted WT or Nupr1$^{fl/fl}$Mx1-cre HSCs were extracted and reverse transcript into cDNA.

(C) Cell cycle analysis of Nupr1$^{fl/fl}$Mx1-cre HSCs under homeostasis. Representative plots of cell cycle from representative WT and Nupr1$^{fl/fl}$Mx1-cre mice (8-week-old).

WT littermates (8-week-old) were used as control. HSCs (Lin$^-$ (CD2$^-$ CD3$^-$ CD4$^-$ CD8$^-$ B220$^-$ Gr1$^-$ CD11b$^-$ Ter119$^-$) CD48$^-$ Sca1$^+$ c-kit$^+$ CD150$^+$ CD34$^-$ CD135$^-$) were analyzed by DNA content (DAPI) versus Ki-67. G0 (Ki-67$^{low}$DAPI$^{2N}$), G1 (Ki-67$^{high}$DAPI$^{2N}$), G2-S-M (Ki-67$^{high}$DAPI$^{>2N-4N}$).

(D) Statistical analysis of the number of LT-HSCs from WT and Nupr1$^{fl/fl}$Mx1-cre mice.

(E) Statistical analysis of HSC cell cycle. **p<0.01; Ctr: n=4, Nupr1$^{fl/fl}$Mx1-cre, n=5.

(F) Kinetic analysis of donor chimerism (CD45.2$^+$) in peripheral blood. Data are analyzed by unpaired Student’s t-test. ***p < 0.001. Data are represented as mean ± SD (Ctr group: n = 5 mice, Nupr1$^{fl/fl}$Mx1-cre group: n =7 mice).

(G) Flow cytometry analysis of HSC compartment in primary recipients four months after transplantation.

(H) Statistical analysis of donor HSC number in the transplantation chimeras. Data are analyzed by unpaired Student’s t-test. Data are represented as mean ± SD **p<0.01, n=5.

### References


1. Yang L, Bryder D, Adolfsson J, et al. Identification of Lin(-)Sca1(+)kit(+)CD34(+)Flt3-short-term hematopoietic stem cells capable of rapidly reconstituting and rescuing myeloablated transplant recipients. Blood. 2005;105(7):2717-2723.
2. Osawa M, Hanada K, Hamada H, Nakauchi H. Long-term lymphohematopoietic reconstitution by a single CD34-low/negative hematopoietic stem cell. Science. 1996;273(5272):242-245.
3. Challen GA, Boles N, Lin KK, Goodell MA. Mouse hematopoietic stem cell identification and analysis. Cytometry A. 2009;75(1):14-24.
4. Kiel MJ, Radice GL, Morrison SJ. Lack of evidence that hematopoietic stem cells depend on N-cadherin-mediated adhesion to osteoblasts for their maintenance. Cell Stem Cell.

2007;1(2):204-217.

5. Tang F, Barbacioru C, Nordman E, et al. RNA-Seq analysis to capture the transcriptome landscape of a single cell. Nat Protoc. 2010;5(3):516-535.

6. Kim D, Langmead B, Salzberg SL. HISAT: a fast spliced aligner with low memory requirements. Nat Methods. 2015;12(4):357-360.

7. Liao Y, Smyth GK, Shi W. featureCounts: an efficient general purpose program for assigning sequence reads to genomic features. Bioinformatics. 2014;30(7):923-930.

8. Love MI, Huber W, Anders S. Moderated estimation of fold change and dispersion for RNA-seq data with DESeq2. Genome Biol. 2014;15(12):550.

9. Subramanian A, Tamayo P, Mootha VK, et al. Gene set enrichment analysis: a knowledge-based approach for interpreting genome-wide expression profiles. Proc Natl Acad Sci U S A. 2005;102(43):15545-15550.
